# Breast Cancer Histopathological Image Classification: A Deep Learning Approach

**DOI:** 10.1101/242818

**Authors:** Mehdi Habibzadeh Motlagh, Mahboobeh Jannesari, HamidReza Aboulkheyr, Pegah Khosravi, Olivier Elemento, Mehdi Totonchi, Iman Hajirasouliha

## Abstract

Breast cancer remains the most common type of cancer and the leading cause of cancer-induced mortality among women with 2.4 million new cases diagnosed and 523,000 deaths per year. Historically, a diagnosis has been initially performed using clinical screening followed by histopathological analysis. Automated classification of cancers using histopathological images is a chciteallenging task of accurate detection of tumor sub-types. This process could be facilitated by machine learning approaches, which may be more reliable and economical compared to conventional methods.

To prove this principle, we applied fine-tuned pre-trained deep neural networks. To test the approach we first classify different cancer types using 6, 402 tissue micro-arrays (TMAs) training samples. Our framework accurately detected on average 99.8% of the four cancer types including breast, bladder, lung and lymphoma using the ResNet V1 50 pre-trained model. Then, for classification of breast cancer sub-types this approach was applied to 7,909 images from the BreakHis database. In the next step, ResNet V1 152 classified benign and malignant breast cancers with an accuracy of 98.7%. In addition, ResNet V1 50 and ResNet V1 152 categorized either benign- (adenosis, fibroadenoma, phyllodes tumor, and tubular adenoma) or malignant- (ductal carcinoma, lobular carcinoma, mucinous carcinoma, and papillary carcinoma) sub-types with 94.8% and 96.4% accuracy, respectively. The confusion matrices revealed high sensitivity values of 1, 0.995 and 0.993 for cancer types, as well as malignant- and benign sub-types respectively. The areas under the curve (AUC) scores were 0.996,0.973 and 0.996 for cancer types, malignant and benign sub-types, respectively. Overall, our results show negligible false negative (on average 3.7 samples) and false positive (on average 2 samples) results among different models. Availability: Source codes, guidelines and data sets are temporarily available on google drive upon request before moving to a permanent GitHub repository.

## I. INTRODUCTION

Recent global cancer statistics reported that breast cancer is still the most common cancer type and the leading cause of cancer-induced mortality among women, worldwide, with 2.4 million new cases and 523,000 deaths per year [19]. Histopathological classification of breast carcinoma is typically based on the diversity of the morphological features of the tumors, comprising 20 major tumor types and 18 minor sub-types ([37]). Approximately 70–80 percent of all breast cancers belongs to either one of the two major histopathological classes, namely invasive ductal carcinoma (IDC) or invasive lobular carcinoma (ILC) ([39], [57]). The IDC class is divided into five different carcinoma sub-types including tubular, medullary, papillary, mucinous and cribriform carcinomas, while benign types of breast cancer contains adenosis, fibroadenoma, phyllodes tumor and tubular adenoma. More importantly, identification of minor tumor sub-types known as special tumor types provides clinically useful information to determine an effective therapy. For instance, accurate diagnosis of tubular and cribriform breast carcinoma can lead to employment of an appropriate treatment and increased overall survival rate. ([57], [12]). A wide range of clinical studies reported lack of complete overlap between immunohistochemically and molecular classification of breast cancer ([6]). However, in 2011 St Gallen International Expert Consensus validated application of immunohistochemistry for identification breast cancer sub-types ([20]). Because of the extensive heterogenecity in breast cancer, and limited predictive power of the histopathological classification, a comprehensive approach for accurate evaluation of cell morphological features is highly required.

To maximize knowledge of cancer detection and interpretation, pathologists have to study large numbers of tumor tissue slides. Additionally, quantification of different parameters (e.g. mitotic counts, surface area and cell size) and evaluation of immunohistochemical molecular markers can be complicated and time consuming. Manual inspection methods introduce three inevitable types of error including statistical, distributional and human errors in low magnification images. These problems adversely affect the accuracy of the differential classification in conventional cancer diagnosis. Therefore, an automated and reproducible methodology could tackle the aforementioned obstacles more effectively.

Computer-aided diagnosis (CAD) established methods for robust assessment of medical image-based examination. In this regard, image processing introduced a promising strategy to facilitate tumor grading and staging, while diminishing unnecessary expenses. Conventional image processing and machine learning techniques require extensive pre-processing, segmentation and manual extraction of specific visual features before classification. However, deep learning approaches have exceeded human performance in visual tasks by utilization of automated hierarchical feature extraction and classification by multi layers, which could be applied for cancer diagnosis using tumor tissue slides.

The first application of the image processing on analytical pathology for cancer detection was introduced by True et al. ([55]), and showed the implication of morphological features in diagnostic methods for malignant tumors. They used a series of morphological features including area fraction, shape, size and object counting to detect cell abnormalities. A large body of evidences has been published concerning cancer detection using various image processing and machine learning techniques ([18], [11], [29], [56], [61], [45]). Application of these methods is limited due to manual feature extraction of the features. Deep learning approach offers an automated, accurate and sensitive method to feature extraction from medical images.

In this regard, the Neighboring Ensemble Predictor (NEP) coupled with Constrained Convolutional Neural Network (SC-CNN) could lead to nucleus detection in colon cancer ([45]). Moreover, AggNet system which is a combination of CNN and additional crowd-sourcing layer, successfully detected mitosis in breast cancer images ([1]). In agreement with this, four deep learning network architectures including GoogLeNet, AlexNet, VGG16 deep network ([58]) and ConvNet with 3, 4, and 6 layers ([13]) were recently applied to identify breast cancer. The best example of using automated CAD system is a study conducted by Esteva and colleague on skin cancer detection using Inception V3, which was done to classify malignancy status ([18]). In addition to these, studies such as ([8], [34], [2], [33]) also showed that deep learning techniques are continuously being applicable to image-based medical diagnosis and improve the performance compared to traditional machine learning techniques.

Despite improvements in images analysis and interpretation, numerous questions related to the reliability and sensitivity of appropriate pathological diagnosis systems particularly for breast cancer classification remained to be answered. In particular, there were no significant, comprehensive and promising solutions for discrimination of breast cancer sub-types.

This study presents deep learning Inception and ResNet architectures to discriminate microscopic cancerous imaging. We demonstrate a highly accurate automatic framework for cancer detection and classification of its sub-types. Our framework also employs additional techniques for data augmentation and advanced pre-processing.

## II. Approach

In this study, we developed and introduced an accurate and reliable computer-based techniques empowered with deep learning approaches to classify cancer types and breast cancer sub-types from histopathological images derived from Hematoxylin and eosin stain (H&E) and Immunohistochemistry (IHC) slides.

Our framework contains five steps: a) Image acquisition and conversion to JPEG/RGB channels. b) Data augmentation (section III-C). c) Deep learning pre-processing (section III-D). d) Transfer learning and fine-tuning pre-trained models (section III-E). e) Hierarchical feature extraction and classification with Inception and ResNet networks (sections III-F). All steps have been illustrated in figure 1.

**Fig. 1.**
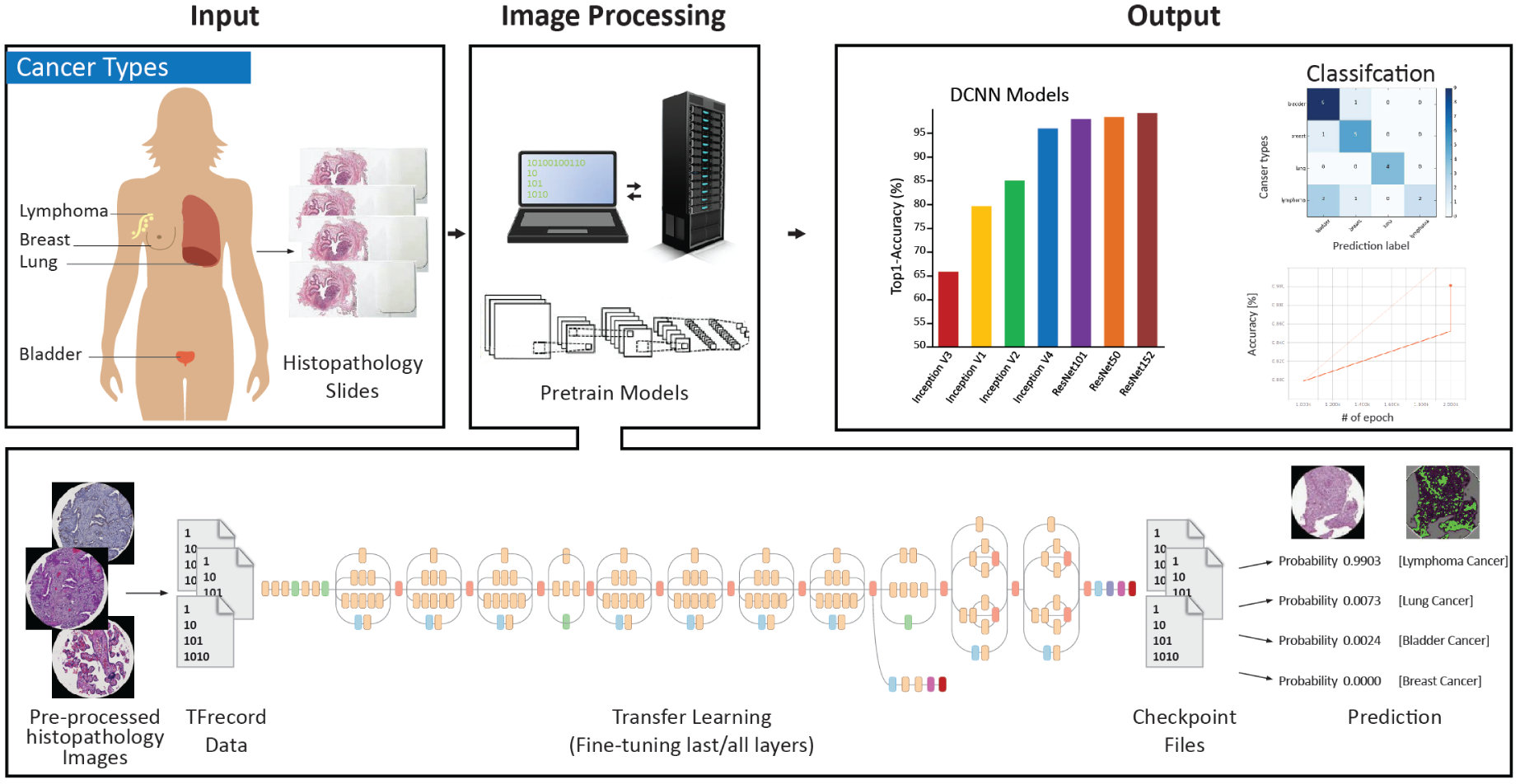
Study work-flow; Data gathering, Image capturing, and Deep learning approaches using per-trained models. Data gathering included captured image from two individual TMA and BreakHis sources saved in JPEG format. Preprocessing is the next step followed by deep learning techniques to extract unique presentation for each separated cancer input.

## III. Materials and Methods

### A. Data-sets

Data-sets were collected from two sources of cancer types and breast cancer sub-types including Tissue Micro Array (TMA) database ([23]) and BreaKHis (The Breast Cancer Histopathological Images) ([48]). 6,402 TMA histopathological images were applied across lung, breast, lymphoma, and bladder cancer tissues. BreaKHis 7,909 pathological breast cancer images (2,480 benign and 5,429 malignant images, each with different magnification of 40X, 100X, 200X, and 400X) from 82 patients were selected for sub-types classification. Our data-set contained four distinct histological sub-types of benign breast tumors: adenosis, fibroadenoma, phyllodes tumor, and tubular adenona; as well as four malignant tumors: ductal carcinoma, lobular carcinoma, mucinous carcinoma and papillary carcinoma (Table I). It should be note that approximately 85% of available data were randomly chosen to construct the learning set. The remaining 15% of the data were used for performance evaluation.

**TABLE I.** Number of images in each class

| Classes | Sub-classes | Training | Testing |
| --- | --- | --- | --- |
| Cancers (6,402) | breast (1,670), bladder (1,870), lymphoma (1,560), lung (1,302) | 5,502 | 900 |
| Breast cancer (7,909) | benign (2,480), malignant (5,429) | 7,100 | 809 |
| Malignant (5,429) | ductal-carcinoma (3,451), lobular-carcinoma (626), mucinous-carcinoma (792), papillary-carcinoma (560) | 4,879 | 550 |
| Malignant Augmented (9,394) | ductal-carcinoma (3,451), lobular-carcinoma (1,881), mucinous-carcinoma (2,379), papillary-carcinoma (1,683) | 8,394 | 1,000 |
| Benign (2,480) | adenosis(444), fibroadenoma (1,014), phyllodes-tumor (453), tubular-adenoma (569) | 2,220 | 260 |
| Benign Augmented (7,452) | adenosis (1,335), fibroadenoma (3,045), phyllodes-tumor (1,362), tubular-adenoma (1,710) | 6,652 | 800 |

### B. Color-map Selection

In this work, we used RGB color-map to preserve tissue structures and different features of histophatological images.

### C. Data Augmentation

Data augmentation is essential step to have enough diverse samples and learn a deep network from the images. Several studies investigated the role of data augmentation in deep learning ([44], [59], [16]). We considered data augmentation for breast cancer sub-types due to difference in the number of images among different sub-type classes. Technically, data augmentation was accomplished on data acquired from Augmentor Python library ([5] see supplementary material) and included random resizing, rotating, cropping, and flipping methods (Figure 2).

**Fig. 2.**
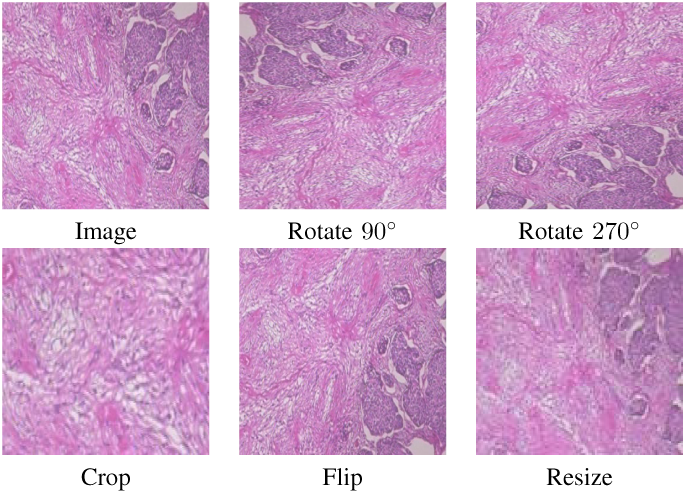
Data augmentation techniques including rotating, cropping, flipping, resizing. Augmentor ([5]) rotated input image into 90 and 270 degrees. Then, image flipped top-bottom to right with 0.8 probability. Next image was cropped with probability of 1 and percentage area of 0.5. Finally, input image was resized with width=120 and height=120

### D. Pre-Processing Steps

Color map selection and data augmentation were followed by pre-processing steps as a preliminary recommended phase to prepare data for further feature extraction and analysis.

Previous studies proposed different pre-processing methods because of the nature of their data ([38], [10], [35], [32], [31]). This work proposed a series of calculation, divided into five steps. The first step focused on JPEG file decoder, followed by TFRecord ([21]) format conversion based on Protocol Buffers ([17], [22], [52]). In third step, TFRecords were normalized to [0,1]. Afterwards, whole image bounding box were re-sized to 299 × 299 × 3 or 224 × 224 × 3 according to the recommended model image size for Inception and ResNet architectures ([50], [49], [26]). Finally, as Inception and ResNet pre-processing, input training images were randomly flipped left to right horizontally and then cropped to create image summaries to display the different transformations on images. In order to improve power of learning and to make the network invariant to aspects of the image that do not affect the label, color distortion with permutation of four hue, brightness, saturation and contrast adjustment operations were applied. On the other hand, in the evaluation step, all images were normalized, cropped and re-sized to specific height and width (Figure 3).

**Fig. 3.**
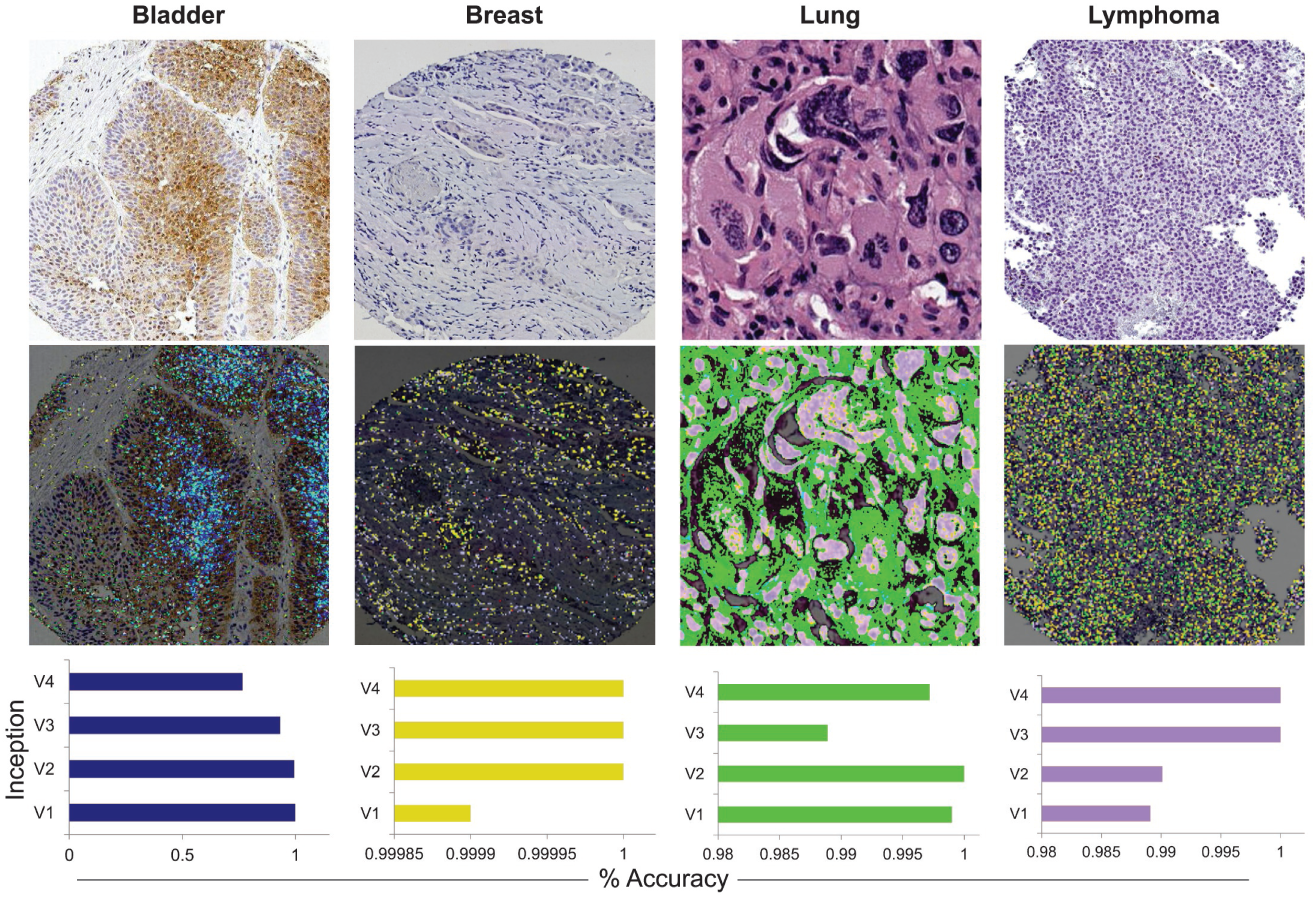
From top to bottom: histopathological images from four cancer types were used as input data. Preprocessing techniques were applied to extract precise learned features. Accuracy of Inception V1 to V4 classification was presented as bar plots

### E. Transfer Learning

Transfer learning is defined as exporting knowledge from previously learned source to a target task ([14], [62], [24], [3]). Learning from clinical images from scratch is often not the most practical strategy due to its computational cost, convergence problem ([51]), and insufficient number of high-quality labeled samples. A growing body of experiments have investigated pre-trained models in the presence of limited learning samples ([63], [15], [43]).

Pre-trained ConvNets alongside fine-tuning and transfer learning lead to faster convergence and outperform training from scratch ([54], [51], [60]). Our target data-set (with 6402 cancer type and 7909 breast cancer sub-types histopathological images) is obviously smaller than the used reference data-set (ImageNet; training data with 1.2M ([53]).

Therefore, we initialized weight of different layers of our proposed network by using ImageNet Inception and ResNet pre-trained models. Then, we employed last layer fine-tuning on cancer images data set. Therefore, the ImageNet pre-trained weights were preserved while the last fully connected layer was updated continuously. Since, the cancer data-sets analyzed here are large and very different from ImageNet, the full layer fine-tuning was applied to compare accurately classification of cancers with the last layer fine tuning([53]) (Table S1)

### F. Inception and ResNet Architectures

Among various deep learning methods, we considered Inceptions and ResNet architectures. It is well understood that Inception models migrated from fully- to sparsely-connected architectures. In order to add more non-linearity capability, Inception module technically included 1 × 1 factorized convolutional neural networks followed by the rectified linear unit (ReLU). Also, a 3×3 convolutional layer was employed. Auxiliary logits with a combination of average pool, convolutional 1 × 1, fully connected, and softmax activation was applied to preserve the low-level detail features and tackle vanishing gradient problem in last layers. ResNet permanently utilized shortcut connections between shallow and deep networks to control and adjust training error rate ([49]).

This study examined different frameworks of Inception (V1, V2, V3, and V4) and ResNet (V1 50, V1 101, and V1 152) ([50], [49], [26]) on cancer digital images. Furthermore, RMSProp adaptive learning rate ([27], [42]) was applied with start- (0.001), decay- (0.9), and end-points (0.0001) settings. Because of insufficient number of available histopathological cancer images (section III-A) compared to numerous model parameters (up to 5 million in Inception and 10 million in ResNet), dropout regularization and batch normalization ([28]) were applied with batch sizes of 32 in training and 100 in evaluation steps.

### G. Computerized System Configuration

Deep learning training with extreme number of network parameters, computational tasks and large data-sets was significantly accelerated by a single computing platform with following specifications: model: HP DL380 G9, CPU: 2x E5-2690v4 (35 MB L3 Cache, 2.6 GHz, 14C), RAM: 64 GB (8 × 8 GB) RAM DDR4 2133 MHz, HDD: 146 GB HDD 7.2k, GPU: ASUS GeForce GTX 1080, 1733 MHz, 2560 CUDA Cores, 8GB GDDR5 with CentOS 7.2 64-bit operating system and Python 3.5.3. In addition, The GPU-enabled version of TensorFlow required CUDA 8.0 Toolkit and cuDNN v5.1 ([36], [21]). All GPU necessary settings and details were obtained from TensorFlow and TFslim documentations and NVIDIA GPUs support ([21]).

## IV. Results

The results were divided into following parts. a) Cancer types classification. b) Cancers were categorized as malignant and benign types. c) Malignant and benign samples were classified into their related four sub-types (sub-section III-A).

Several standard performance terms such as true positive (TP), false positive (FP), true negative (TN), false negative (FN), accuracy (ACC), precision (P), AUC and sensitivity (S) were isolated from the confusion matrix ([46]).

### A. Classification of Cancer Types

A 4 × 4 confusion matrix was used to represent prediction results of the set of four cancer pathological samples (subsection III-A). The matrices were built on four rows and four columns: breast, lung, bladder, and lymphoma representing the known cancer classes. Statistical performance measurement of each cancer type and different deep learning frameworks (section III-F) were summarized in Tables II and III. The result indicated that ResNet V1 50 and fine-tuning all layers classified 99.8% of known cancer types. While this rate decreased to 99.6% for ResNet V1 101/152 fine-tuning all layers with 3,000 epochs (Table III). ResNet V1 101 with 3,000 epochs and fine-tuning last layer had an accuracy rate of 99.5% (Table II). The ResNet models showed significantly increased accuracy for four cancer type classification compared to the all Inception structures, whereas Inception V1 with 3,000 epochs and fine-tuning all layer showed 97.1% accuracy at best. Additionally, there was an obvious difference in false positive values between Inception structures and ResNets. Furthermore, on average less false positive results were obtained by the ResNet (0.3) in comparison to the Inception models (82) with 3, 000 epochs (Table III). The Cohens unweighted kappa coefficient statistic ([40]) of the Inception V1, V2, V3 and V4 fine-tuning all layers with 3,000 epochs were 0.94,0.94,0.82 and 0.84 respectively,while those of the ResNet models were above 0.97. The Inception networks were able to correctly identify four types of cancer with an accuracy ranging between 76.4% and 97.1%, compared to the ResNet networks with ranging between 98.3% and 99.8% (Table III, 3,000 epochs).

**TABLE II.** Fine-tuning the last layer for different models in cancer type Classification

| Model name | Epochs | ACC | TP | TN | FP | FN | P | AUC | S |
| --- | --- | --- | --- | --- | --- | --- | --- | --- | --- |
| Inception V1 | 3,000 | 0.917 | 580 | 252 | 14 | 54 | 0.976 | 0.917 | 0.914 |
| Inception V2 | 3,000 | 0.848 | 516 | 252 | 14 | 118 | 0.973 | 0.874 | 0.813 |
| Inception V3 | 3,000 | 0.884 | 568 | 244 | 22 | 66 | 0.962 | 0.869 | 0.895 |
| Inception V4 | 3,000 | 0.871 | 542 | 258 | 8 | 92 | 0.985 | 0.905 | 0.854 |
| ResNet V1 50 | 3,000 | 0.993 | 630 | 266 | 0 | 4 | 1 | 0.996 | 0.993 |
| ResNet V1 101 | 3,000 | 0.995 | 625 | 272 | 1 | 1 | 0.998 | 0.994 | 0.998 |
| ResNet V1 152 | 3,000 | 0.992 | 623 | 272 | 1 | 4 | 0.998 | 0.993 | 0.993 |

**TABLE III.**
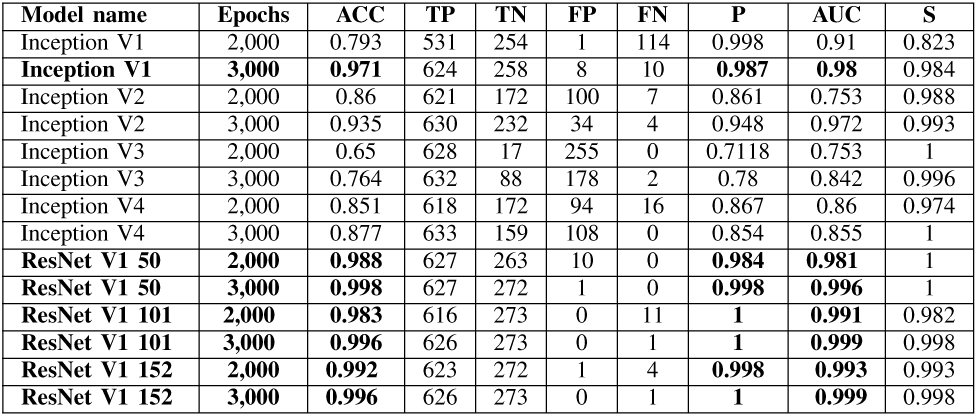
Fine-tuning all layers for different models in cancer type Classification

### B. Malignant and Benign Breast Cancer

The breast cancer data (section III-A and Table I) were categorized into malignant and benign groups. Using an 90% training set and 10% test set,the ResNet V1 101 fine-tuning all layers, correctly classified malignant and benign cancer types with 98.4% confidence (Table IV). This performance for ResNet V1 50 with fine-tuning all layers and ResNet V1 101 with fine-tuning the last layer decreased to 97.8% and 94.1% respectively (Tables S2 and IV).

**TABLE IV.**
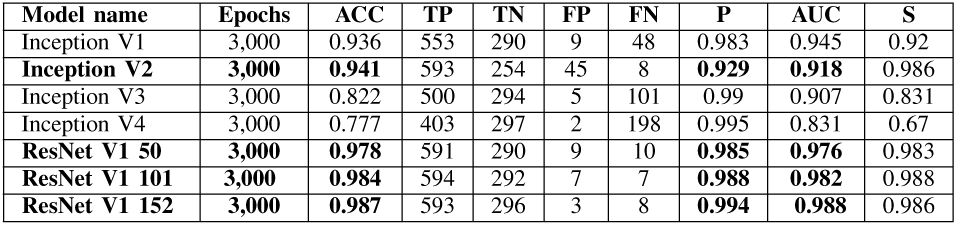
Fine-tuning all layers for different models in breast cancer classification

| Model name | Epochs | ACC | TP | TN | FP | FN | P | AUC | S |
| --- | --- | --- | --- | --- | --- | --- | --- | --- | --- |
| Inception V1 | 3,000 | 0.936 | 553 | 290 | 9 | 48 | 0.983 | 0.945 | 0.92 |
| **Inception V2** | **3,000** | **0.941** | 593 | 254 | 45 | 8 | **0.929** | **0.918** | 0.986 |
| Inception V3 | 3,000 | 0.822 | 500 | 294 | 5 | 101 | 0.99 | 0.907 | 0.831 |
| Inception V4 | 3,000 | 0.777 | 403 | 297 | 2 | 198 | 0.995 | 0.831 | 0.67 |
| **ResNet V1 50** | **3,000** | **0.978** | 591 | 290 | 9 | 10 | **0.985** | **0.976** | 0.983 |
| **ResNet V1 101** | **3,000** | **0.984** | 594 | 292 | 7 | 7 | **0.988** | **0.982** | 0.988 |
| **ResNet V1 152** | **3,000** | **0.987** | 593 | 296 | 3 | 8 | **0.994** | **0.988** | 0.986 |

Inception V2 with fine-tuning all layers, indicated the maximum accuracy (94.1%) among the Inception architectures (Table IV). Moreover, the results showed on average less false positive in the ResNet (6.3) compared to the Inception models (15.25) (Table IV).

### C. Breast Cancer Sub-types Classification

In order to create a framework for approval classification capability, we considered wide varieties of similar and complex histopathological images related to different sub-types of breast cancer. Since the benign and malignant groups were well separated from each other (section IV-B), further, we assessed pre-trained Inception and ResNet models to classify benign and malignant related sub-types. According to our results, the accuracy of analysis for benign sub-types resulted to classification of adenosis, fibroadenoma, phyllodes-tumor, and tubular-adenoma (Tables S3 and V).

**TABLE V.** Fine-tuning all layers for different models in augmented benign data.

| Model name | Epochs | ACC | TP | TN | FP | FN | P | AUC | S |
| --- | --- | --- | --- | --- | --- | --- | --- | --- | --- |
| Inception V1 | 3,000 | 0.696 | 660 | 88 | 48 | 4 | 0.932 | 0.779 | 0.993 |
| **Inception V2** | **3,000** | **0.723** | 660 | 95 | 41 | 4 | **0.941** | **0.875** | 0.993 |
| Inception V3 | 3,000 | 0.512 | 638 | 44 | 92 | 26 | 0.873 | 0.629 | 0.96 |
| Inception V4 | 3,000 | 0.54 | 651 | 53 | 83 | 13 | 0.886 | 0.404 | 0.98 |
| **ResNet V1 50** | **3,000** | **0.948** | 661 | 132 | 4 | 3 | **0.993** | **0.973** | 0.995 |
| **ResNet V1 101** | **3,000** | **0.933** | 658 | 132 | 4 | 6 | **0.993** | **0.971** | 0.99 |
| **ResNet V1 152** | **3,000** | **0.945** | 662 | 129 | 7 | 2 | **0.989** | **0.98** | 0.996 |
| **ResNet V1 101** | **6,000** | **0.965** | 659 | 134 | 2 | 5 | **0.996** | **0.992** | 0.992 |

In case of malignant classification, accuracy analysis for associated sub-types (on test sets) resulted in 96.4%, and 94.6% for ResNet V1 152 and ResNet V1 50 with fine-tuning all layers respectively (Table VII). Moreover, an accuracy rate of 90% for ResNet V1 152 with fine-tuning the last layer is also acceptable (Table VI).

**TABLE VI.** Fine-tuning the last layer for different models in augmented malignant data

| Model name | Epochs | ACC | TP | TN | FP | FN | P | AUC | S |
| --- | --- | --- | --- | --- | --- | --- | --- | --- | --- |
| Inception V1 | 3,000 | **0.794** | 631 | 366 | 1 | 2 | **0.998** | **0.997** | 0.996 |
| Inception V2 | 3,000 | 0.647 | 456 | 366 | 1 | 177 | 0.997 | 0.857 | 0.72 |
| Inception V3 | 3,000 | 0.74 | 632 | 365 | 2 | 1 | 0.996 | 0.995 | 0.998 |
| Inception V4 | 3,000 | 0.745 | 624 | 362 | 5 | 9 | 0.992 | 0.981 | 0.985 |
| ResNet V1 50 | 3,000 | **0.893** | 632 | 367 | 0 | 18 | **1** | **0.999** | 0.998 |
| ResNet V1 101 | 3,000 | **0.872** | 630 | 367 | 0 | 3 | **1** | **0.997** | 0.995 |
| ResNet V1 152 | 3,000 | **0.9** | 627 | 367 | 1 | 5 | **0.998** | **0.995** | 0.992 |

**TABLE VII.** Fine-tuning all layers for different models in augmented malignant data

| Model name | Epochs | ACC | TP | TN | FP | FN | P | AUC | S |
| --- | --- | --- | --- | --- | --- | --- | --- | --- | --- |
| Inception V1 | 3,000 | **0.866** | 631 | 368 | 0 | 1 | **1** | **0.999** | 0.998 |
| Inception V2 | 3,000 | 0.565 | 283 | 366 | 1 | 350 | 0.996 | 0.721 | 0.447 |
| Inception V3 | 3,000 | 0.739 | 633 | 367 | 0 | 0 | 1 | 1 | 1 |
| Inception V4 | 3,000 | 0.799 | 634 | 364 | 2 | 0 | 0.996 | 0.999 | 1 |
| ResNet V1 50 | 3,000 | **0.946** | 624 | 367 | 1 | 8 | **0.998** | **0.993** | 0.987 |
| ResNet V1 101 | 3,000 | **0.844** | 529 | 367 | 0 | 104 | **1** | **0.917** | 0.835 |
| ResNet V1 152 | 3,000 | **0.964** | 629 | 367 | 0 | 4 | **1** | **0.996** | 0.993 |
| ResNet V1 101 | 6,000 | **0.985** | 633 | 367 | 0 | 0 | **1** | **1** | 1 |

As represented in (Table VII), the ResNet networks output illustrated significantly higher level of accuracy than other Inception structures in which Inception V1 with 3, 000 epochs and fine-tuning all layers showed 86.6% marked as less accurate method in terms of malignant cancer sub-types classification. Additionally, there has been an obvious difference in false positive values between Inception structures and ResNets with average of 0.75 and 0.33, respectively (Table VII).

A 4 × 4 confusion matrix was used to represent different possibilities of the set of instances. The matrices represented distribution of the ductal-carcinoma, lobular-carcinoma, mucinous-carcinoma, papillary-carcinoma in different classes. It was well evidenced that increases in the number of epochs could improve the accuracy of classification. Our findings (Tables V and VII) suggest that ResNet with higher epochs was highly accurate (for example; 96.5% and 98.5% ResNet 101 with 6, 000 epochs) for classification of specific sub-types of breast cancer (Figure 4).

**Fig. 4.**
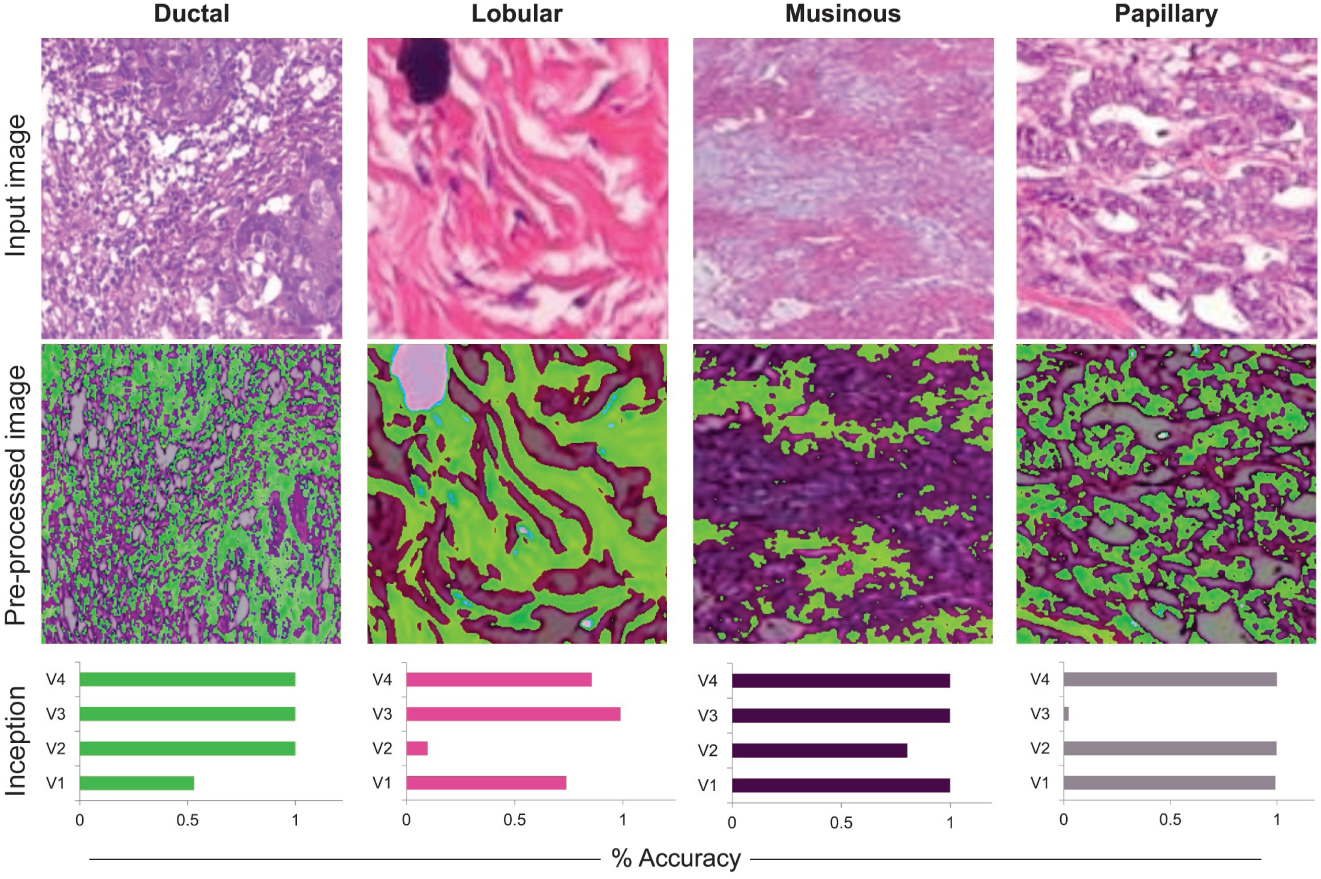
From top to bottom: histopathological images from four cancer types were used as input data. Preprocessing techniques were applied to extract precise learned features. Accuracy of Inception V1 to V4 classification was presented as bar plots

## V. Discussion

Breast cancer multi-classification aim to identify sub-classes of breast cancer (adenosis, fibroadenoma, phyllodes tumor, tubular adenoma, ductal carcinoma, lobular carcinoma, mucinous carcinoma, and papillary carcinoma) among samples found in broad variability of resolution image appearances, high coherency of cancerous cells, and extensive inhomogeneity of color distribution. This work examined data gathered from TMA and BreaKHis data-sets (see section III) to classify cancer types and breast sub-types. Previous studies ([30], [47], [48]) focused on binary benign-malignant classification and did not perform further quantitative assessment. In this work, we introduced automated breast cancer multi-classification methods. We suggested a generic CAD framework based on deep networks for learning histopathology images to avoid hand-crafted pathological features. In this study, we compared the performance of Inception and ResNet deep learning models using transfer learning strategy on several large image data-sets. We found that deep ResNet models were more sensitive and reliable than Inception in all tested cancer data-sets. We combined different magnification including 40X, 100X, 200X and 400X to generate comprehensive, independent and scalable system while a large number of previous studies employed single magnification level ([7], [30]). Several other studies ([7], [25], [48], [4]) also investigated multiple magnifications of medical images. However, these approaches examined different classifiers for each magnification level and also had medical laboratory limitations to capture required multiple magnification to gather image training samples.

In recent comparative studies, ([48], [30], [7]), conventional machine learning (SVM, KNN, QDA, ASSVM, SSVM-SCAD, etc) along with hand-crafted feature extraction were used. The results were evaluated at various magnifications (i.e. 40X, 100X, 200X and 400X). To data, the benign and malignant classification had significantly high accuracy rates ranging from 90 to 93%. In addition, to classify benign and malignant, AlexNet deep learning approach with numerous learning parameters resulted in an accuracy rate of 90% ([47]). Moreover, Han and colleagues (Han et al., 2017), reported a deep learning-based multi classification of breast cancer with an average accuracy rate of the 93.2%.

In conclusion, the ResNet frameworks with 99.8%, 98.7%, 94.8%, and 96.4% accuracy for four cancer types, two main breast cancer types, benign and malignant related sub-types and trivial false positive average values (0.3 out of 900 for four cancer types, 6.3 out of 809 for all breast cancer, 5 out of 800 for benign and 0.3 out of 1000 for malignant) were able to examine histopathological images obtained by different imaging devices with different magnification levels. This multi-classification system relieves the pathologists and medical experts workloads regarding to analyze and interpretation of the histopathological slides for assisting the doctors to choose more efficient therapeutic approaches.

## VI. Conclusion and Future Work

Using deep learning ResNet approach with specific settings for cancer detection is an effective and reliable strategy compared to the conventional approaches. This work concentrated on the application of the proposed frameworks to cancer subtypes detection. In this research, various Inception and ResNet deep learning classifications are presented and the use of these theories is outlined. The findings are expected to be more comprehensively evaluated and discussed by future works considering different deep learning semantic segmentation algorithms (i.e., U-net: convolutional networks for biomedical image segmentation ([41]) and DeepLab v3 ([9])). The empirical findings of this study provided a better understanding of deep learning in medical applications. The framework principals could extend in the field of pathological analysis and computer-assisted diagnosis using medical images.

## VII. Acknowledgment

We would like to acknowledge Dr. Mahdi Pourfath from the University of Tehran and his team for a wide range of resources on GPU programming. We would like to appreciate comments and ideas by Dr. Ali Sharifi Zarchi.

